# Enhancer-promoter competition between homologous alleles leads to reduced transcription in early *Drosophila* embryos

**DOI:** 10.1101/2021.08.16.456541

**Authors:** Hao Deng, Bomyi Lim

**Affiliations:** Department of Chemical and Biomolecular Engineering, University of Pennsylvania, Philadelphia, Pennsylvania, United States

## Abstract

The mechanism by which transcriptional machinery is recruited to enhancers and promoters to regulate gene expression is one of the most challenging and extensively studied questions in modern biology. Here, we ask if inter-allelic interactions between two homologous alleles can affect gene regulation. Using MS2- and PP7-based, allele-specific live imaging assay, we visualized de novo transcription of a reporter gene in hemizygous and homozygous *Drosophila* embryos. Surprisingly, each homozygous allele produced fewer RNAs than the hemizygous allele, suggesting the possibility of allelic competition in homozygotes. Moreover, the *MS2-yellow* reporter gene showed reduced transcriptional activity when a partial transcription unit (enhancer or promoter only) was in the homologous position. We propose that the transcriptional machinery that binds to both the enhancer and promoter region, such as RNA Pol II or preinitiation complexes, may be responsible for the allelic competition. To support this idea, we showed that the homologous alleles did not interfere with each other in earlier nuclear cycles when Pol II is in excess, while the degree of interference gradually increased in nuclear cycle 14. Such allelic competition was observed for endogenous *snail* as well. Our study provides new insights into the role of 3D inter-allelic interactions in gene regulation.

## Introduction

Enhancers, which contain multiple binding sites for sequence-specific transcription activators and repressors, determine when and where a target gene should be transcribed (Levine et al., 2014; Levine & Tjian, 2003). Missense mutations in enhancers or disruptions in enhancer-promoter interactions often result in ectopic or lost expression of target genes (Halder et al., 1995; Riedel-Kruse et al., 2007). Many of these genetic perturbations in enhancers are known to be associated with various disease phenotypes (Hnisz et al., 2013; Miguel-Escalada et al., 2015). Hence, precise control of enhancer-promoter communications plays a major role in the normal development of multicellular eukaryotes.

Extensive studies have been conducted to elucidate the mechanism of enhancer-mediated transcriptional regulation, and yet, there still remain more questions to be answered. For example, multiple enhancers that drive target gene expression in the same tissue need to coordinate among themselves to ensure proper access to the target promoter (Berrocal et al., 2020; Lim, Fukaya, et al., 2018; Perry et al., 2011). A live imaging study on early *Drosophila* embryos demonstrated that some enhancers work additively with each other, while others work sub-additively or even super-additively, such that multiple enhancers can drive significantly higher or lower transcription activity than a single enhancer (Bothma et al., 2015). Although it is still not understood how multiple enhancers interact simultaneously with the target promoter, this result suggests that multivariate interactions among transcription units affect the extent of RNA production.

Meanwhile, a single enhancer can interact with multiple promoters as well. A study in early *Drosophila* embryos showed that a single enhancer can co-activate two *cis-*linked reporter genes, rather than driving exclusive transcription of a single promoter (Fukaya et al., 2016). In the presence of insulators, an enhancer on one allele can also interact in *trans* with the target promoter on the homologous allele to initiate transcription, a phenomenon known as transvection (Lim, Heist, et al., 2018). Moreover, this *trans-*activation is always accompanied by a co-activation of the *cis-*linked reporter gene, indicating that a single shared enhancer can activate both the *cis-* and the *trans-*linked reporter genes (Fukaya et al., 2016; Lim, Heist, et al., 2018). These recent studies suggest that enhancer-promoter communication is more dynamic than previously thought.

In parallel, multiple recent studies have shown that various transcriptional regulators form local clusters at active transcription loci (Chong et al., 2018; Kato et al., 2012; Sabari et al., 2018). These clusters include RNA Pol II, Mediators, pre-initiation complexes (PICs), and site-specific TFs (Cho et al., 2016; Cisse et al., 2013; Wollman et al., 2017). It was suggested that TF clusters at enhancers and Pol II/Mediator clusters at promoters form an active hub to regulate transcription (Boija et al., 2018; Tsai et al., 2017). Indeed, several studies in *Drosophila* embryos showed that local highly concentrated clusters of the pioneer factor Zelda (Zld) at transcription loci facilitate the binding of Bicoid (Bcd) and Dorsal (Dl) activators to the target loci (Mir et al., 2017, 2018; Yamada et al., 2019). This “transcription hub” idea can also explain previous findings on multivariate enhancer-promoter interactions where one enhancer can co-activate two target promoters (Fukaya et al., 2016; Lim, Heist, et al., 2018). Altogether, these studies propose that dynamic enhancer-promoter interactions and clustering of transcriptional machinery in a nucleus play important roles in gene regulation.

In this study, we provide evidence that two homologous alleles may compete with each other to affect the level of RNA production. Using allele-specific MS2- and PP7-based live imaging methods in early *Drosophila* embryos, we measured the transcription activity of one allele from homozygous and hemizygous embryos containing a reporter gene. Surprisingly, we found that a hemizygous allele produced more RNA than a single homozygous allele. The change in transcriptional activity was manifested mainly as a change in transcriptional amplitude, implying that the number of RNA Pol II loaded to the promoter was reduced in homozygotes. Interestingly, this allelic competition at the homologous locus was observed only in the presence of strong enhancer-promoter interactions. When we replaced a strong enhancer with a weaker one, or weakened the enhancer-promoter interactions by placing a strong enhancer 6.5 kb away from the promoter, we did not observe a reduction in transcriptional activity of the homozygous allele.

To examine whether shared transcriptional machinery between the two alleles could explain the reduced RNA production from the homozygotes, we measured the transcriptional activity of a reporter gene under two conditions: enhancer only and promoter-reporter gene only in the homologous locus. Unexpectedly, this partial transcription unit on the homologous position was sufficient to decrease the reporter gene’s transcriptional activity. This implies that the transcriptional machinery binding to both the enhancer and promoter can cause competition between alleles. Based on these results, we propose that homologous alleles may share the same local transcription hub, and that each allele produces a reduced number of RNAs when the number of Pol II in the hub is limiting – especially upon strong enhancer-promoter interactions. Indeed, we showed that the competition was observed only in the nuclear cycle 14 (NC14) when massive zygotic genome activation occurs. Lastly, we demonstrated that endogenous *snail* alleles also interfere with each other. We believe that our study provides new insights into a mechanism of transcriptional regulation in 3D environments.

## Results

### MS2- and PP7-based labeling of two homologous alleles

In order to test the possibility that homologous alleles may interact with each other, we compared the transcriptional activity of reporter genes driven by the well-characterized *snail* shadow enhancer (snaSE) between hemizygous and homozygous embryos (Figure 1A) (Perry et al., 2010). MS2- and PP7-based live imaging methods, which were successfully implemented in *Drosophila* embryos and many other tissues, were used to visualize nascent transcripts (Chen et al., 2018; Coulon et al., 2013, 2014; Fukaya et al., 2017; Larson et al., 2011). Twenty-four copies of the MS2 or PP7 sequences were inserted directly upstream of the *yellow* reporter gene (Figure 1B). Upon transcription, MS2 or PP7 sequences form a stem-loop structure, each of which can be recognized by two copies of the MS2 coat protein (MCP) or the PP7 coat protein (PCP), fused with GFP and mCherry, respectively. The binding of MCP-GFP or PCP-mCherry to the transcribed MS2 or PP7 stem loops allows visualization of de novo transcriptional activity in living embryos (Figure 1C) (Lim, Heist, et al., 2018). We generated transgenic lines where snaSE and the 100-bp core promoter of *sna* drive expression of the *MS2-yellow* and the *PP7-yellow* reporter gene. Reporter genes were inserted into a specific location in the 3^rd^ chromosome using PhiC31-mediated site-directed transgenesis (Bischof et al., 2007; Groth, 2004).

**Figure 1.**
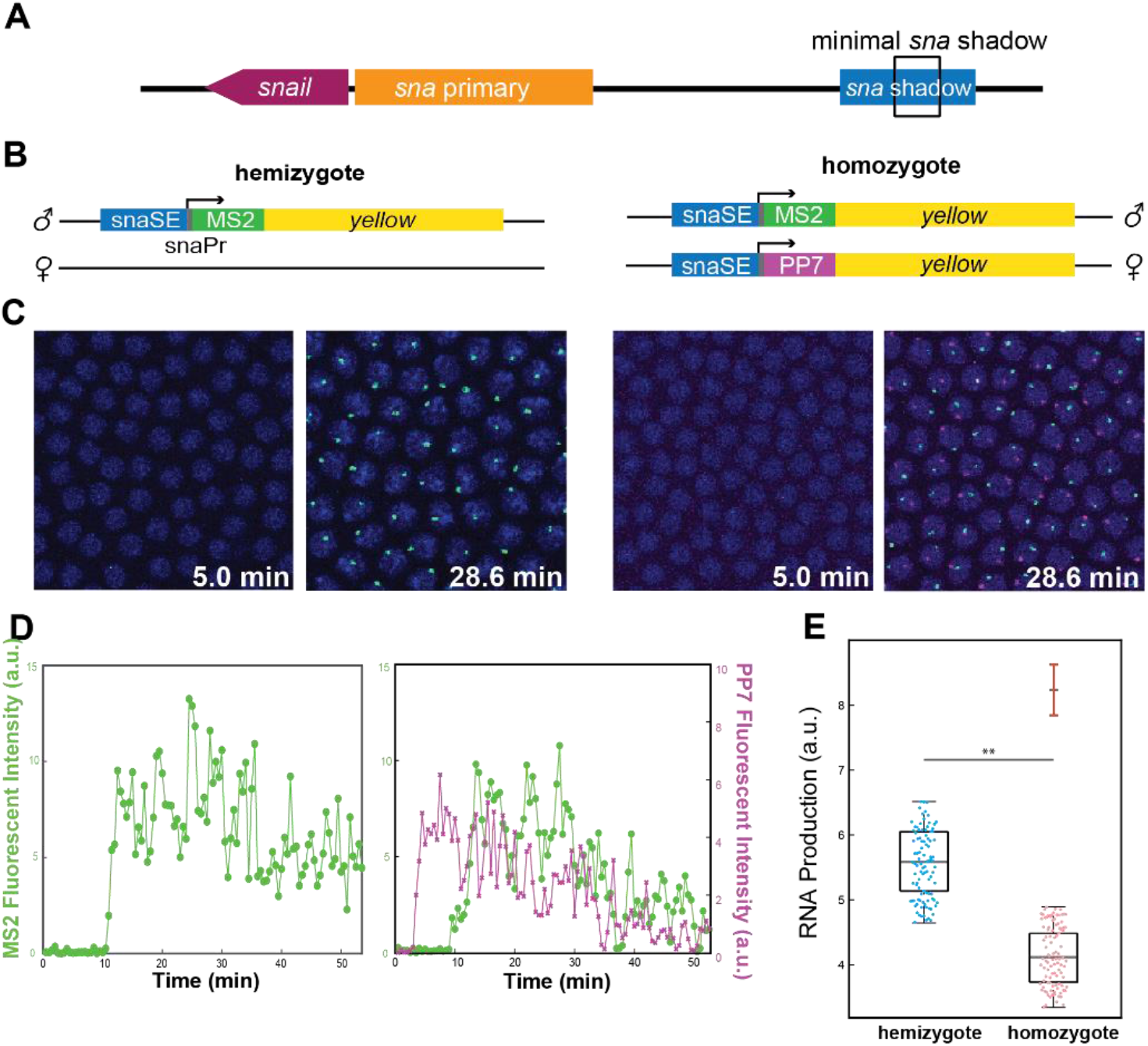
Allelic competition between the homologous alleles reduces transcriptional activity. (A) Schematic of the *snail* gene (*sna*), the primary enhancer (snaPE), the shadow enhancer (snaSE), and the minimal shadow enhancer (snaSEmin). The snaSEmin is a minimal version of snaSE that contains fewer transcription factor binding sites. (B) Schematic of the hemizygous and homozygous snaSE>*yellow* constructs. snaSE is placed right upstream of the *sna* promoter. 24 copies of MS2 and PP7 sequences were inserted for allele-specific visualization of the transcriptional activity. The paternal allele contains the *MS2-yellow* reporter gene for both hemizygotes and homozygotes. (C) Representative snapshots of hemizygous and homozygous embryos containing the snaSE>*yellow* transgene. The time indicates minutes after the onset of NC14. Fluorescence puncta (green - *MS2-yellow;* red - *PP7-yellow*) indicate active nascent transcripts. Nuclei were visualized with His2Av-eBFP2 (blue). (D) Representative transcriptional trajectories from a nucleus of hemizygous and homozygous embryos shown in (C). Transcriptional activity is proportional to the fluorescence intensity. The total RNA production of each allele can be estimated by measuring the area under the transcription trajectory. (E) Boxplot showing RNA production of the snaSE>*MS2-yellow* allele in hemizygous and homozygous embryos. The scatter points show RNA production of two hundred random nuclei from the analysis. The boxplots show that the *MS2-yellow* allele from homozygotes produced approximately 25% fewer RNAs than the one from hemizygotes. The mark above the homozygote boxplot shows the projected total RNA production of a homozygous embryo, which is obtained by doubling the RNA production of one allele in homozygotes. Hemizygotes produce around 70% of the total RNA production of homozygotes. 1,060 and 2,908 nuclei from 4 and 10 biologically replicate embryos were analyzed for hemizygotes and homozygotes, respectively. The box indicates the 25%, 50%, and 75% quantile, and the whiskers extend to the 10th and 90th percentile of each distribution. ** indicates p<1E-4.

To distinguish transcriptional activities from each allele in homozygous embryos, we crossed MCP-GFP, PCP-mCherry / snaSE*>PP7-yellow* females with snaSE*>MS2-yellow* / snaSE>*MS2-yellow* males. 50% of the progeny have two copies of the *yellow* reporter gene, each marked with *PP7* and *MS2* stem-loops (homozygous embryos). The other 50% have one copy of the *yellow* reporter gene marked with MS2 stem-loops (hemizygous embryos) (Figure 1C and Movie 1). To note, the paternal allele carries the *MS2-yellow* reporter gene for both homozygous and hemizygous embryos. In order to analyze if the alleles from homozygous embryos behave independently of each other, we compared the transcriptional activity of the *MS2-yellow* allele between the homozygous and the hemizygous embryos.

### Live imaging reveals a possibility of allelic competition between the homologous alleles

Since our live imaging methods provide instantaneous transcriptional activity as a function of time, we can estimate total RNA production by measuring the area under the transcriptional trajectory of each nucleus (Figure 1D). Theoretically, the two alleles from a homozygous embryo should produce equal amounts of RNAs. In other words, if the alleles do not interact in *trans*, a single allele from homozygous embryos should produce a comparable amount of RNAs as the allele from hemizygous embryos. In the case of competition between the two homologous alleles, the transcriptional activity of each homozygous allele would differ from that of the hemizygous allele. To our surprise, the *MS2-yellow* allele in homozygous embryos produced about 25% fewer RNAs than the *MS2-yellow* allele in hemizygous embryos (Figure 1E).

If we assume that both maternal and paternal alleles from homozygous embryos produce similar amounts of RNA, the total RNA output from homozygous embryos could be estimated to be twice that from the single *MS2-yellow* allele. This indicates that the homozygous alleles combined make less than twice the RNA produced by the single hemizygous allele (Figure 1E). We hypothesize that the homologous alleles may interact in *trans* and inhibit each other, resulting in transcription of fewer RNAs than expected.

We acknowledge that if there is a bias in RNA production between the maternal and the paternal alleles, the alleles may complement each other instead of interfering with each other. To examine this possibility, we generated two homozygous snaSE>*MS2-yellow /* snaSE>*PP7-yellow* embryos, one with the maternal *MS2-yellow* reporter gene and the other with the paternal *MS2-yellow* reporter gene. We confirmed that the *MS2-yellow* transcriptional activity does not change between maternal and paternal alleles (Figure S1). This indicates that the 25% reduction in RNA production observed in homozygous embryos applies to both maternal and paternal alleles, resulting in ~1.5 times more RNA production in homozygotes compared to hemizygotes (rather than twice more). In sum, our allele-specific live imaging assays suggest that the homologous alleles of snaSE>*yellow* reporter genes seem to interfere with each other and result in significantly lower transcriptional activity (Figure 1E).

### Homozygous alleles produce fewer RNA mainly due to lower transcriptional amplitude

We investigated the source of this reduced RNA production by taking advantage of our single-cell resolution live imaging data. There are three possible scenarios that could result in decreased RNA output: (i) late onset of transcription, (ii) less time spent in the transcriptionally active state, and (iii) reduction in transcriptional amplitude. The onset of transcription could be delayed if there was a lag in enhancer-promoter interactions to initiate transcription. Alternatively, if Pol II is loaded less frequently to the promoter, this could result in less frequent transcriptional bursting, hence a shorter duration of active transcription and reduced RNA production. Lastly, a reduction in the number of Pol II loaded to the promoter could lead to a decrease in transcriptional amplitude.

To distinguish these factors, we measured 1) the timing of transcription initiation, 2) the duration of active transcription, and 3) the average amplitude of transcription in each transcriptionally active nucleus. We found that transcription was initiated about 6 min after the onset of NC14 in both hemizygous and homozygous embryos (Figure 2A). The duration of active transcription was also comparable between the two genotypes, with about 5% shorter duration for the homozygous allele (Figure 2B). Note that we used the duration of active transcription as a proxy for the frequency of transcriptional bursting since fewer bursting events would lead to a shorter transcriptionally active state. Unlike these two parameters that showed minimal effect, the average amplitude of transcriptional activity was significantly different between the two genotypes, where the homozygous *MS2-yellow* allele exhibited about 20% lower amplitude than the hemizygous *MS2-yellow* allele (Figure 2C). Indeed, when we examined the average trajectory for the homozygous and the hemizygous *MS2-yellow* allele, the amplitude was the main difference between the two genotypes (Figure 2D).

**Figure 2.**
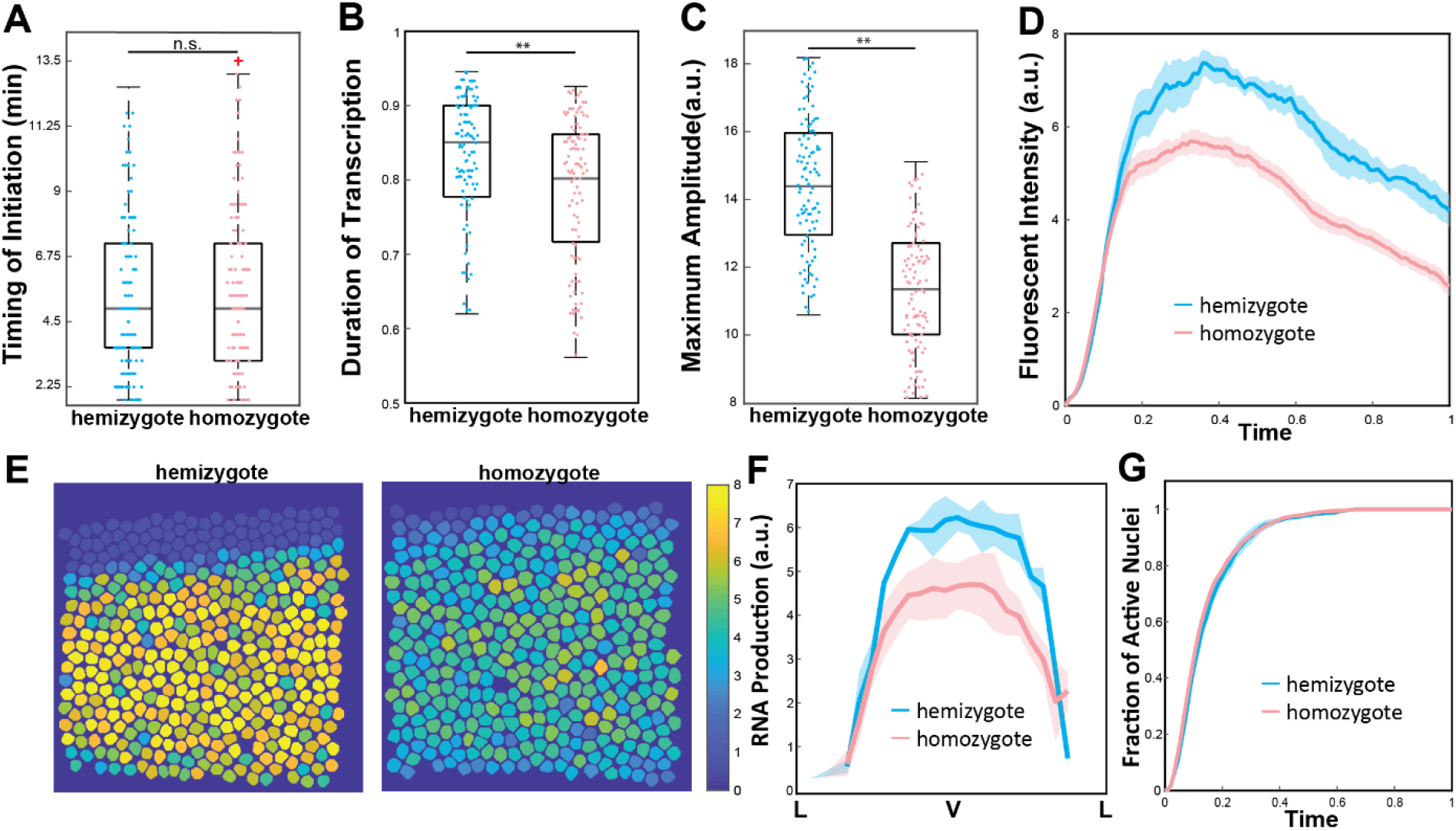
Low transcriptional amplitude caused reduced RNA output in the homozygous alleles. (A) Boxplot of the timing of transcription initiation for hemizygotes and homozygotes expressing snaSE>*yellow*. For both genotypes, transcription was initiated about 6 min after the onset of NC14. (B) Boxplot of the duration of active snaSE>*MS2-yellow* transcription in NC14. The hemizygous and the homozygous alleles spend comparable time in the active transcription state. (C) Boxplot of the average amplitude of *MS2-yellow* fluorescenct intensity in snaSE>*yellow* hemizygotes and homozygotes. The amplitude in homozygous embryos is about 20% lower than the one in hemizygous embryos. (D) Average transcriptional trajectories of active nuclei from hemizygous (blue) and homozygous (red) embryos. The main difference between the genotypes is the average amplitude. (E) Heat maps of a representative snaSE>*yellow* hemizygous (left) and homozygous (right) embryo showing the accumulated RNA production in NC14 of all nuclei within the *sna* expression domain. The RNA production is reduced throughout the ventral side of the homozygous embryo. The snapshot shows a ventral view of an embryo. (F) Average RNA production of hemizygotes (blue) and homozygotes (red) expressing the snaSE>*MS2-yellow* reporter gene along the dorsoventral axis of an embryo. The RNA production is reduced throughout the domain in homozygotes yet the *sna* expression boundary is not narrowed. (G) Plot of the cumulative fraction of active nuclei over the duration of NC14 in hemizygotes (blue) and homozygotes (red). Both genotypes produce RNAs with similar kinetics of transcriptional activation. The number of analyzed nuclei is the same as the one shown in Figure 1E. For boxplots in (A-C), the scatter points indicate values from 200 randomly selected nuclei used in the analysis. The box indicates the 25%, 50%, and 75% quantile, and the whiskers extend to the 10th and the 90th percentile of each distribution. The error bar in (D), (F), and (G) represents the Standard Error of Mean (SEM) for 4 and 10 biologically replicate embryos for hemizygous and homozygous snaSE>*yellow* embryos, respectively. ** indicates p<1E-4.

In addition to the single-cell analysis, we analyzed if all the nuclei within the *sna* expression domain were affected uniformly, or if the boundary nuclei were affected to a greater extent. *sna* is expressed in the ventral side of an embryo with an expression of 18-20 cell width. Its sharp domain establishes a boundary between the presumptive mesoderm and neurogenic ectoderm (Ip et al., 1992; Kosman et al., 1991). The *MS2-yellow* allele from homozygous embryos demonstrated reduced RNA production throughout the *sna* expression domain, without narrowing the expression domain or affecting the spatial pattern along the dorsoventral axis (Figure 2E and F). We also measured the cumulative fraction of active nuclei over time, and both the homozygous and the hemizygous *MS2-yellow* allele exhibited similar kinetics of transcriptional activation (Figure 2G). This result indicates that the rate of forming the *sna* expression boundary is about the same between the two genotypes. Similar to what we observed in the single-cell analysis, the level of RNA production decreased, but the overall width and pattern of the *sna* boundary were maintained (Figure 2F). These results suggest that the alleles may compete in *trans* throughout the *sna* expression domain, mainly by modulating transcriptional amplitude.

### Weaker enhancer-promoter interactions do not result in allelic interference

We wondered if the decreased transcription activity in homozygous snaSE>*yellow* alleles is valid for other enhancers as well. Transgenic lines with the minimal *sna* shadow enhancer (snaSEmin) were generated to address this. The snaSEmin encompasses 520bp within the *sna* shadow enhancer with fewer transcription factor binding sites for Dorsal (Dl) or Twist (Twi) activators (Ferraro et al., 2016). The snaSEmin drives transcription only from late NC13, while the snaSE induces transcription as early as NC10. Despite the delay in transcription, the expression pattern driven by the minimal enhancer is similar to the one driven by the shadow enhancer (Ferraro et al., 2016).

Much like the shadow enhancer constructs, we made transgenic lines where *MS2-yellow* and *PP7-yellow* reporter genes were regulated by the snaSEmin (Figure 3A). Crosses were made such that homozygous embryos carry the paternal *MS2-yellow* allele and maternal *PP7-yellow* allele. Hemizygous embryos have only the *MS2-yellow* reporter gene in their paternal allele (Figure 3A). We compared the RNA production of the *MS2-yellow* allele between hemizygous and homozygous embryos. Unlike what was observed in the snaSE constructs, this time there was no evidence of allelic competition as both genotypes produced similar amounts of RNAs (Figure 3D). Subsequently, all other parameters of transcription activity were comparable between hemizygotes and homozygotes (Figure S2).

**Figure 3.**
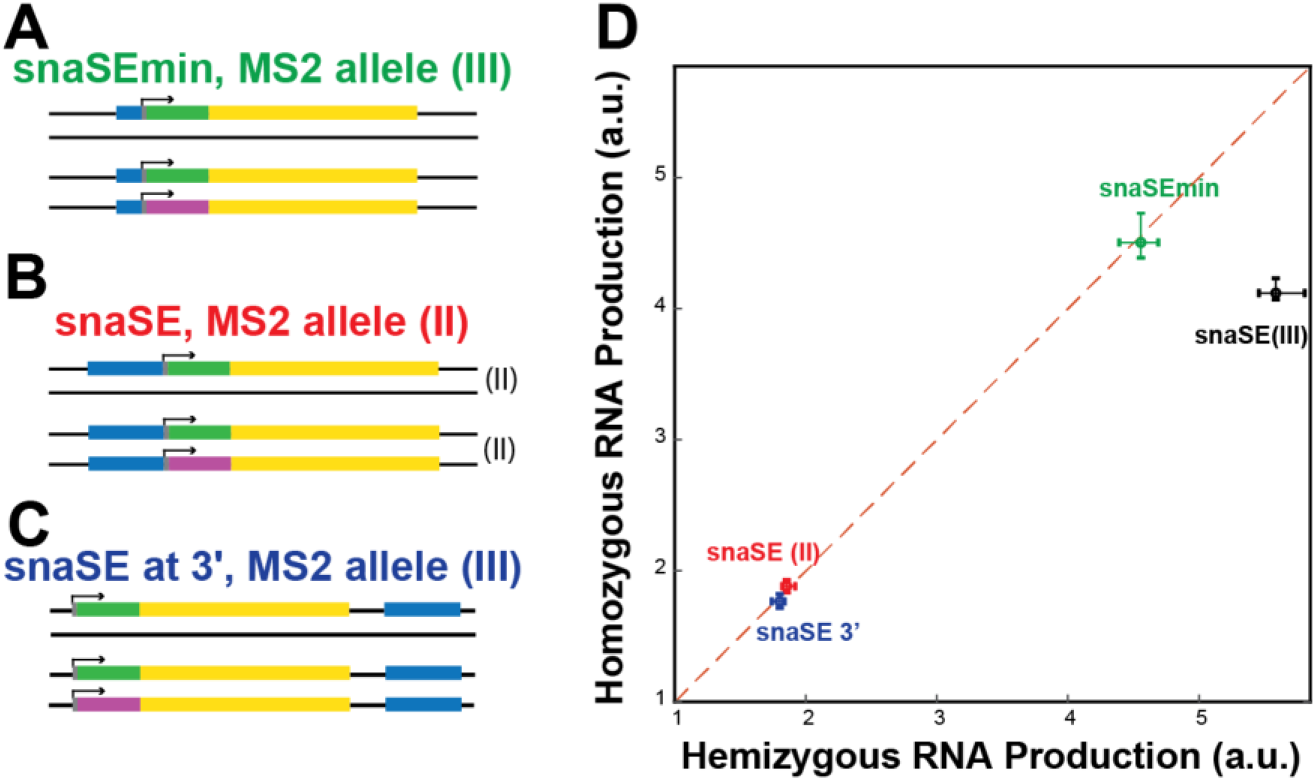
Allelic competition is not observed when the enhancer-promoter interactions are weakened. (A-C) Schematic of hemizygous and homozygous constructs with (A) snaSEmin>*yellow* reporter genes in the 3^rd^ chromosome, (B) snaSE>*yellow* reporter genes in the 2^nd^ chromosome, and (C) snaSE>*yellow* reporter genes in the 3^rd^ chromosome, where the snaSE enhancer (blue) is inserted downstream of the reporter gene. (D) Plots comparing the RNA production of the *MS2-yellow* allele between the homozygous and the hemizygous constructs shown in (A)-(C) and snaSE>*yellow* reporter genes in the 3rd chromosome. In all three constructs shown in (A)-(C), the RNA production is lower than the one from the snaSE>*yellow* constructs shown in Figure 1, and the alleles do not compete with each other. The error bars represent 99% confidence intervals. 1,334 and 1,210 nuclei from 4 and 4 biologically replicate snaSEmin>*yellow* hemizygous and homozygous embryos were analyzed. 599 and 756 nuclei from 4 and 4 biologically replicate snaSE>*yellow* (II) hemizygous and homozygous embryos were analyzed. 293 and 271 nuclei from 4 and 4 biologically replicate snaSE3′>*yellow* hemizygous and homozygous embryos were analyzed.

We wondered if weakened enhancer-promoter interactions between the snaSEmin and the target promoter could affect the degree of allelic interactions. To test this idea, we inserted the strong snaSE>*yellow* constructs, which showed allelic competition, in different chromosomal contexts. The original *sna*SE-driven reporter gene constructs were inserted into the 3^rd^ chromosome using the PhiC31-mediated site-directed transgenesis (VK00033 line) (K. J. T. Venken et al., 2006; Koen J T Venken et al., 2009). It was reported that the same transgene could produce different amounts of RNAs depending on the chromosomal contexts, due to differences in histone methylation/acetylation levels, and TADs boundaries, etc. (Levis et al., 1985; Wallrath & Elgin, 1995). Therefore, we inserted the same transgene into the 2^nd^ chromosome instead, using the VK00002 line (Koen J T Venken et al., 2009). Despite the use of the same strong snaSE, transcriptional activity was reduced by about 65% compared to the 3rd chromosome insertion (Figure 3D). Surprisingly, the *sna*SE>*MS2-yellow* that exhibited allelic interference in the 3^rd^ chromosome location did not show competition in the 2^nd^ chromosome location and produced a similar amount of RNAs (Figure 3D).

We created another transgenic line with the snaSE by inserting the snaSE downstream of the reporter gene (Figure 3C). It was previously shown that a larger distance between an enhancer and the target promoter leads to lower transcriptional activity (Fukaya et al., 2016; Yokoshi et al., 2020). The transgene was inserted into the original VK00033 position in the 3^rd^ chromosome. As expected, the amplitude of transcription was lower when the enhancer was inserted downstream of the reporter gene (Figure 3D). The two *MS2-yellow* alleles from the hemizygous and homozygous embryos were comparable, indicating that the two homozygous alleles behave independently of each other (Figure 3D). Since we used the strong *sna*SE enhancer in the same 3^rd^ chromosome position, there should be no difference in the number of TFs bound to the enhancer. However, the 6.5kb distance between the enhancer and the promoter weakened the enhancer-promoter interaction. Our results suggest that allelic interference seems to be associated with the level of enhancer-promoter interactions, such that alleles with stronger enhancer-promoter interactions compete with each other while the alleles with weaker interactions do not.

### Alleles with partial transcription units can still compete with each other

We next examined potential mechanisms of the observed allelic competition. One possible explanation is that transcription factors that are available to bind to enhancers are limiting. A recent study showed that a limiting number of transcription factors could lead to reduced RNA production from the homozygous allele (Waymack et al., 2021). Since the snaSEmin has fewer Dl and Twi activator binding sites, fewer transcription factors will be needed for the snaSEmin homozygotes, and hence the number of TF may not be limiting, resulting in no allelic interference. This idea, however, does not fully explain the absence of competition when the snaSE>*MS2-yellow* was inserted into a different chromosome, or when the strong snaSE was placed downstream of the promoter (Figure 3B and C).

Another hypothesis is that the limiting amount of Pol II and other PIC molecules induced the reduction in transcription activity in homozygous embryos. With recent studies on enhancer RNAs, it is known that Pol II binds to both the enhancer and the promoter regions to initiate transcription at both locations (Adelman & Lis, 2012; Kim et al., 2010; Savic et al., 2015). It was also shown that in early *Drosophila* embryos, the amount of TATA-Binding Protein (TBP) and TAFII is limiting (Mannervik, 1999; Zhou et al., 1998). Based on such previous studies, we hypothesized that two homologous alleles may share the same transcription hub where the number of Pol II and PIC factors are limiting, resulting in reduced transcriptional activity (please see Discussion).

Two additional constructs were designed to test this idea. One allele has an intact *sna* shadow enhancer, 100-bp core *sna* promoter, and the *MS2-yellow* reporter gene, while the homologous allele contains either only the *sna* shadow enhancer without the reporter gene (“enhancer only”) or the *even-skipped* (*eve*) promoter-reporter gene cassette without the *cis-*linked enhancer (“promoter only”) (Figure 4A and B). If the amount of transcription factor is limiting, interference will occur for the constructs with the “enhancer only” allele since transcription factors will still bind to the enhancer region on both alleles. On the other hand, the interference should not be observed for the constructs with the “promoter only” allele, as transcription factors do not bind to the core promoter region, and the *MS2-yellow* allele is expected to behave similarly to the hemizygous allele.

**Figure 4.**
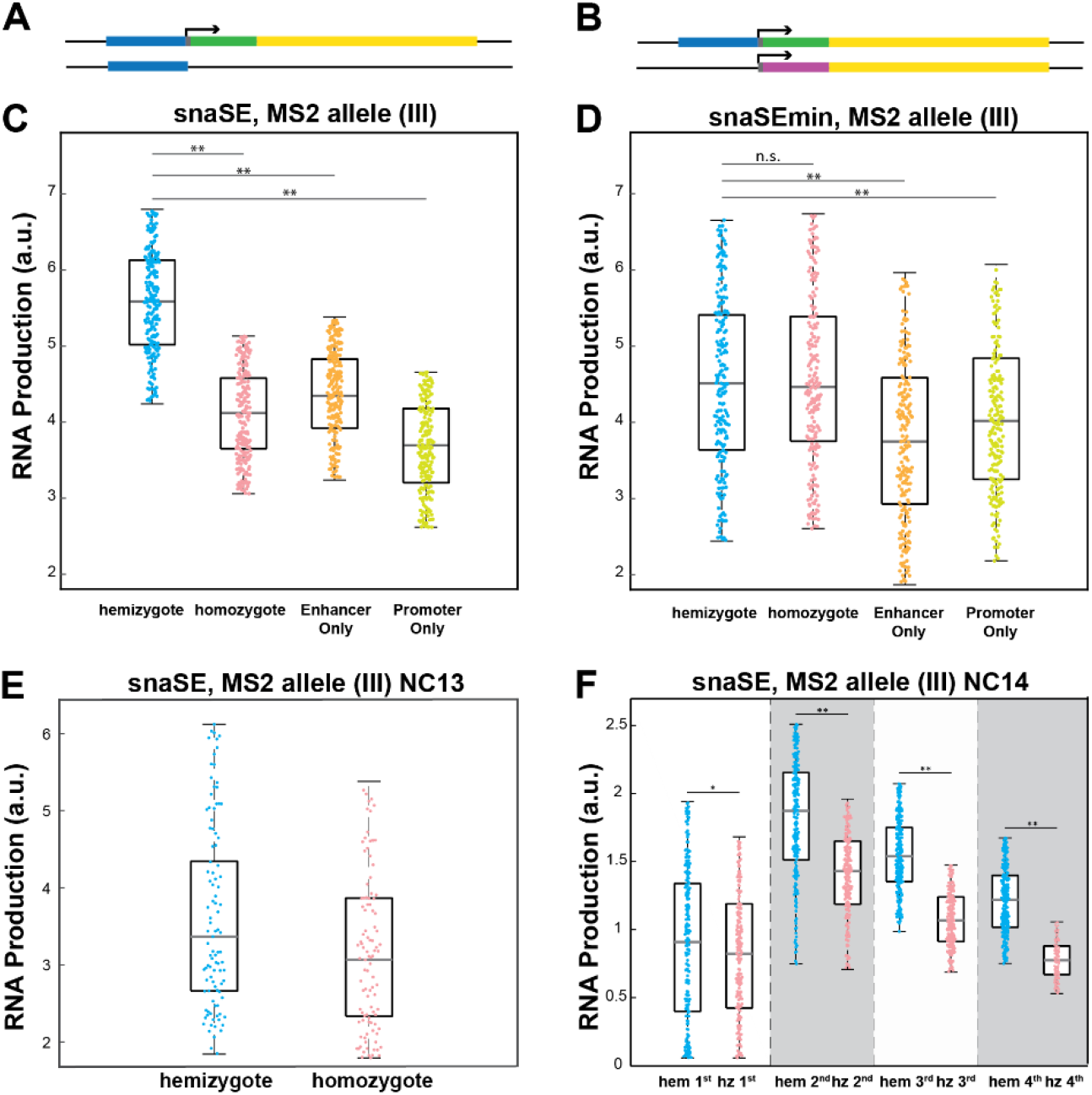
Allelic competition is observed with a partial transcription unit and in the late nuclear cycle. (A-B) Schematics of the (A) “enhancer only” and the (B) “promoter only” constructs. Both constructs contain the enhancer>*MS2-yellow* on the paternal allele. The “enhancer only” construct has the *sna* shadow enhancer, and the “promoter only” construct has the core *eve* promoter and the *PP7-yellow* reporter gene in the homologous position. (C-D) Boxplots showing RNA production of the *MS2-yellow* allele from the hemizygous, homozygous, enhancer only, and promoter only embryos containing (C) snaSE>*yellow* or (D) snaSEmin>*yellow*. For both constructs, the *MS2-yellow* allele exhibits significant reduction in transcriptional activity in the presence of an enhancer only or the *PP7-yellow* reporter gene only on the homologous position. The number of analyzed nuclei for snaSE>*yellow* and snaSEmin>*yellow* hemizygous and homozygous embryos is shown in Figure 1E and Figure 3, respectively. 1,076 nuclei and 1,100 nuclei from 4 and 4 biologically replicate embryos were analyzed for snaSE>*yellow* and snaSEmin>*yellow* “enhancer only” constructs, respectively. 984 nuclei and 1,142 nuclei from 4 and 4 biologically replicate embryos were analyzed for snaSE>*yellow* and snaSEmin>*yellow* “promoter only” constructs, respectively. (E) Boxplot showing RNA production of the snaSE>*MS2-yellow* during NC13. No allelic competition is observed. (F) Boxplot showing the RNA production of the snaSE>*MS2-yellow* during NC14. RNA production was measured in four temporal classes in NC14 by dividing the duration of NC14 into four. In early NC14, transcriptional activities of both genotypes are comparable to each other. Later in NC14, however, a reduced expression is observed in homozygotes with a greater difference towards the end of NC14. For all boxplots, the box indicates the 25%, 50%, and 75% quantile, and the whiskers extend to the 10th and 90th percentile of each distribution. The scattered points indicate values from 100 (E) or 200 (F) randomly selected nuclei used in the analysis. * indicates p<1E-3 and ** indicates p<1E-4.

Surprisingly, we found that the transcriptional activity of the *MS2-yellow* allele from both the “enhancer only” and the “promoter only” constructs behaved like the *MS2-yellow* allele from homozygous constructs, exhibiting reduced transcriptional activity compared to the hemizygotes (Figure 4C). What was more unexpected was that this reduction in transcriptional activity was observed for the constructs with the minimal *sna* shadow enhancer as well (Figure 4D). The snaSEmin>*MS2-yellow* allele from hemizygous and homozygous embryos exhibited similar transcriptional activity as shown in Figure 3. However, a decrease in transcriptional activity was observed when the snaSEmin>*MS2-yellow* allele was in homologous position as the “enhancer only (*sna* shadow)” or the “promoter only” allele (Figure 4D). There was about a 15% decrease in transcriptional activity for snaSEmin>MS2-yellow, compared to the ~25% reduction observed for snaSE>MS2-yellow constructs. While we have yet to investigate the cause of the interference between the intact allele and the “enhancer only” or the “promoter only” allele, our results show that the alleles can interfere with each other even if the homologous allele has only a partial transcription unit. Also, this suggests that Pol II and PIC factors, the molecules that bind to both enhancers and promoters, may play a role in allele competition.

### Allelic interference is not observed in early NC but is in late NC

We wanted to further test the idea that the limiting amount of local Pol II and PIC hubs prompts allelic interference. We analyzed transcriptional activity between hemizygous and homozygous embryos at earlier NCs when limited zygotic transcription occurs, and hence fewer Pol II are needed. In NC13, about ~950 zygotic genes are activated, compared to the activation of ~3,500 genes in NC14 (Kwasnieski et al., 2019). We hypothesized that such massive activation of the zygotic genome in NC14 could greatly consume local Pol II and PIC in each hub, leading to reduced transcriptional activity from each allele. In accordance with the hypothesis, we found that during NC13, the transcriptional activity of the *sna*SE>*MS2-yellow* allele was comparable between hemizygous and homozygous embryos (Figure 4E). We then divided NC14 into four temporal classes (0-25%, 25-50%, 50-75%, and 75-100% of NC14) and examined how the RNA production differs between the hemizygous and the homozygous embryos. RNA production was similar between the two genotypes in early NC14. However, the homozygous *MS2-yellow* allele produced fewer RNAs compared to the hemizygous *MS2-yellow* allele, showing bigger differences towards late NC14 (Figure 4F).

The transcriptional machinery is non-uniformly distributed in a limited number of transcription hubs in a given nucleus (Boehning et al., 2018; Edelman & Fraser, 2012; Mir et al., 2017; Tsai et al., 2019; Yamada et al., 2019; Zhu et al., 2021). Even if numerous hubs exist in each nucleus, their positions could be relatively fixed and heterogeneously localized, preventing each hub to move freely towards active transcription loci. Instead, each gene would use the transcriptional machinery in a nearby hub to process transcription. In our case, in the vicinity of the *yellow* reporter gene, there may exist just a single hub at that specific 3D location that is accessible by the reporter gene. Therefore, our observations of allelic competition in NC14 but not in NC13 support the hypothesis that the limiting amount of local Pol II and PICs could lead to the observed allelic competition.

### Endogenous sna also exhibits allelic competition

So far, we have used transgenic reporter genes to provide evidence that the two homologous alleles compete with each other. We wondered if a homozygous allele of endogenous genes also produces fewer RNAs than a heterozygous allele. Using CRISPR-mediated genome editing, we inserted MS2 and PP7 stem loops to the 3′ UTR of the endogenous *sna* to generate *sna-MS2* and *sna-PP7* lines (Figure 5A). By crossing *sna-MS2 / sna-MS2* flies with *sna-PP7 / CyO* flies, we obtained either hemizygous *sna-MS2 / CyO* or homozygous *sna-MS2 / sna-PP7* embryos. We compared the transcriptional activity of the *sna-MS2* allele for homozygous and hemizygous embryos. Similar to the transgenic lines, we found that the endogenous *sna* alleles also compete with each other such that the homozygous *sna-MS2* allele produces fewer RNAs than the hemizygous allele (Figure 5C). Moreover, this interference was only observed in NC14 but not in NC13, agreeing with the results from the snaSE>*MS2-yellow* transgenic lines (Figure 4E and 5B). Our results with endogenous *sna* suggest that the allelic competition may be a general feature of transcriptional regulation for some strongly expressed genes. Taken together, we believe that the localized cluster of Pol II and PICs along with specific transcription factors form “transcription hubs” within a nucleus, capping the total RNA production level for some strong genes and result in reduced transcriptional activity of homozygous alleles.

**Figure 5.**
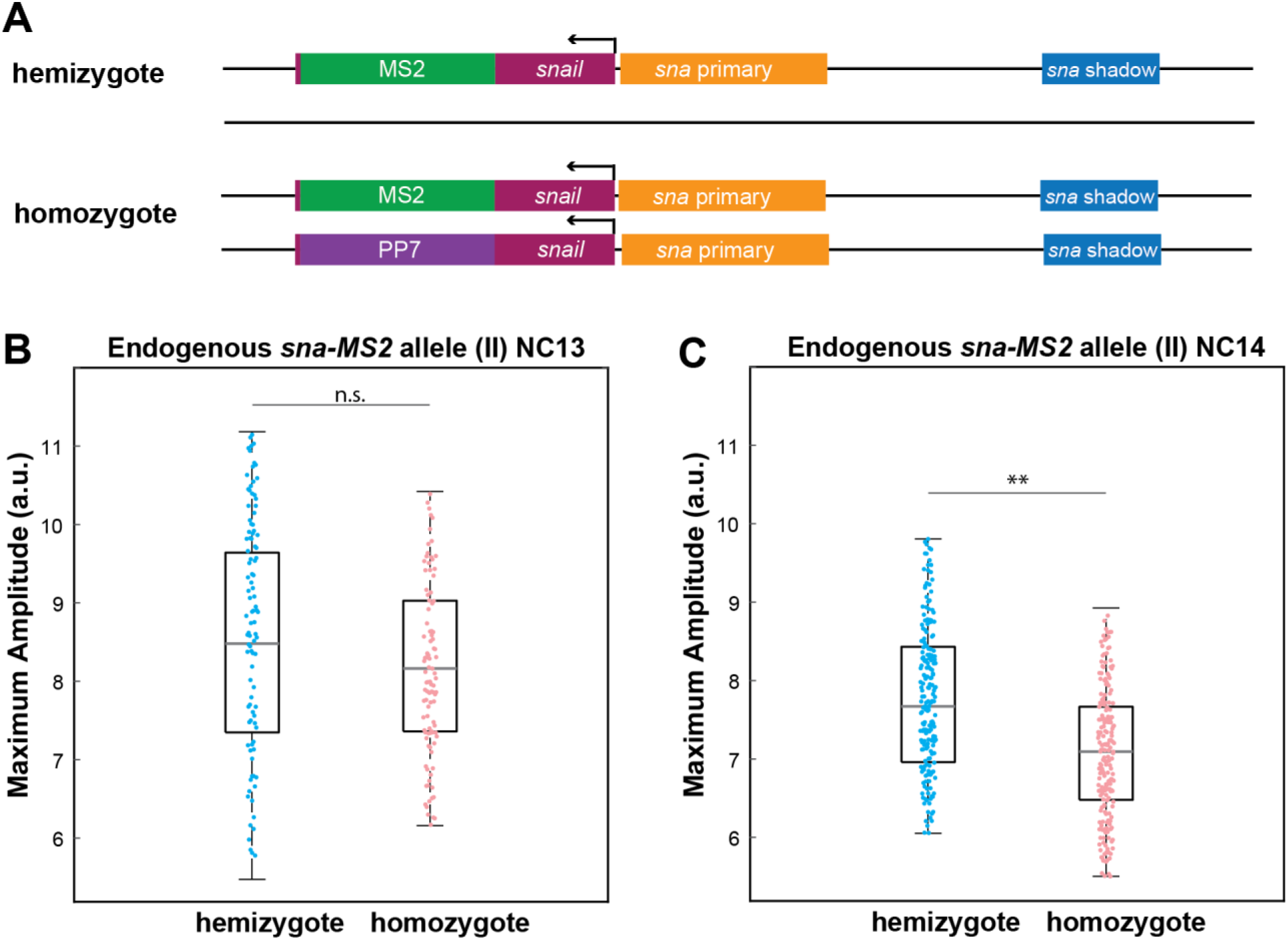
Allelic competition is observed for endogenous *sna*. (A) Schematic of the hemizygous and homozygous endogenous *sna* constructs. MS2 or PP7 stem loops are inserted into the 3′UTR of the endogenous *sna* via CRISPR-mediated genome editing. (B-C) Boxplot of the transcriptional amplitude of the hemizygous and homozygous *sna-MS2* alleles during (B) NC13 and (C) NC14. No allelic competition is observed in NC13. On the other hand, amplitude of the homozygous *sna-MS2* allele is about 10% lower than the hemizygous allele’s amplitude. 293 and 308 nuclei from 2 and 2 biologically replicate NC13 *sna-MS2* hemizygous and homozygous embryos were analyzed. 734 and 564 nuclei from 3 and 3 biologically replicate NC14 *sna-MS2* hemizygous and homozygous embryos were analyzed. For boxplots in (B-C), the scatter points indicate values from 100 (B) or 200 (C) randomly selected nuclei used in the analysis. The box indicates the 25%, 50%, and 75% quantile, and the whiskers extend to the 10th and the 90th percentile of each distribution. ** indicates p<1E-4.

## Discussion

Here, we have shown that homozygous alleles may interfere with each other to produce fewer RNAs per allele than a hemizygous allele. Strikingly, this decrease in RNA production was observed even when the homologous allele contained only a partial transcription unit such as an enhancer or reporter gene only. We have presented evidence to support the hypothesis that the local concentration of Pol II in transcription hubs may be limiting, thus leading to allele competition and reduced transcriptional activity.

A recent study demonstrated a similar reduction of transcriptional activity in homozygotes compared to the hemizygotes, using reporter genes driven by the *Krüppel* enhancers (Waymack et al., 2021). The authors showed that inserting an array of Bcd or Zld TF binding sites on the homologous position was sufficient to reduce the transcription activity of the reporter gene, suggesting that the limiting number of TFs may induce allelic competition. This idea is in agreement with our finding where the stronger *sna* shadow enhancer exhibits allelic interference while the weaker minimal *sna* shadow enhancer does not (Figure 3). Since the minimal enhancer has fewer Dl and Twi binding sites than the full shadow enhancer, the number of TFs may be sufficient, resulting in no allelic competition. However, we also showed that the transcriptional activity from the snaSE>*MS2-yellow* or the snaSEmin>*MS2-yellow* allele was reduced when the homologous allele has only the core *eve* promoter and the *MS2-yellow* reporter gene without the enhancer (Figure 4C and D). Since the site-specific TFs like Dl and Twi do not bind to the 100bp-core promoter region, we do not think that the number of site-specific TFs is the only limiting factor that is responsible for the allelic interference.

Instead, we suggest that general TFs or RNA Pol II levels may be also limiting for strongly activated genes. Previous studies showed that the level of TATA-Binding Protein (TBP) and TAFII is limiting in *Drosophila* (Aoyagi & Wassarman, 2001; Colgan & Manley, 1992). For example, one study in early *Drosophila* embryos showed that in the sensitized Dl heterozygous background, TBP or TAFII deletion in one allele leads to defects in *sna* expression (Zhou et al., 1998). In our study, we demonstrated that homozygous alleles that contain strongly expressed reporter genes produce fewer RNAs than the hemizygous alleles. If the number of PICs indeed works as a rate-limiting factor, available proteins need to be divided between the two homologous alleles to initiate transcription, whereas all of them can bind to the hemizygous allele’s promoter, resulting in a lower transcription level for the homozygotes.

Moreover, we showed that the snaSE>*MS2-yellow* allele competes not only with the homologous snaSE>*PP7-yellow* allele, but also competes with an allele with snaSE only or with the *PP7-yellow* reporter gene only (Figure 4C and D). This suggests that a common factor that binds both to the enhancer and the promoter is responsible for the observed allelic interference. Many recent papers have provided evidence that enhancers are actively transcribed, producing enhancer RNAs (eRNAs) (Arnold et al., 2020; Kim et al., 2015). The eRNAs are produced through the binding of Pol II, Mediators, and other general TFs to the enhancer region (Kim et al., 2010). This indicates that Pol II and PICs bind to both enhancers and promoters to induce transcription. Hence, we suggest that the limiting amount of Pol II or PICs can lead to reduced transcription activity in homozygotes. In support of this hypothesis, the RNA production was comparable between the homozygous and the hemizygous allele in NC13, when fewer genes are transcribed. On the other hand, in NC14, where thousands of genes are activated, the transcription activity from the homozygous allele was lower than the one from the hemizygous allele, and this difference increased as the embryo progressed to late NC14 (Figure 4F). These results support our claim that the number of Pol II and PICs can become limiting in early embryos, and this may affect the allelic competition.

We acknowledge that thousands of genes are being transcribed in early embryos, and it is not intuitive to think that one additional transgene can affect the overall balance of TFs, Pol II, and PICs in each nucleus. At the same time, however, others have reported similar phenomena of allelic competition, and we demonstrated that endogenous *snail* alleles interfere with each other as well (Figure 5) (Waymack et al., 2020, 2021). We suggest that the localized clustering of Pol II and TFs in each nucleus allows allelic competition. In the past few years, many papers have reported the presence of “transcription hubs” where TFs, Mediators, PICs, and Pol II form a cluster and the genes within each hub share the transcriptional machinery (Cho et al., 2018; Cisse et al., 2013; Mir et al., 2017; Yamada et al., 2019). Such clustering of molecules was observed and characterized both in *Drosophila* embryos and in mammalian cells (Boija et al., 2018; Dufourt et al., 2018; Wollman et al., 2017). According to the model, only a handful of each transcriptional machinery exist in a given hub, and adding one more reporter gene may work as a rate-limiting factor in this localized environment. Thermodynamic modeling from a recent paper showed that the local competition for TFs can account for the observed decrease in transcription activity (Waymack et al., 2021). Along the same line, local competition of PICs or Pol II can also explain the allelic interference. Taken together, we suggest that localized clusters of transcription hubs can limit the number of available molecules that bind to the enhancer and promoter regions, inducing allelic competition for strongly expressed genes. Our study provides additional insight into how the distribution of Pol II clusters in the nucleus and subsequent inter-allelic competitions can affect enhancer-mediated transcriptional regulation.

## Materials and Methods

### Generation of transgenic lines

Pbphi vectors containing an attB site and the *MS2-yellow-*αtub 3′ UTR and *PP7-yellow-*αtub 3′UTR reporter genes were used to generate transgenic lines used in this study (Fukaya et al., 2016). The 100bp core *sna* or *eve* promoter was amplified and inserted upstream of the *MS2* sequence. *sna* shadow enhancer was amplified using primers (5′ - GCA TTG AGG TGT TTT GTT - 3′) and (5′ - TAA ATT CCG ATT TTT CTT - 3′). Minimal *sna* shadow enhancer was amplified using primers (5′ - CCT TGG TCC TAC CTT CGA - 3′) and (5′ - CCA AAG GCA ACG CCG ATT - 3′). Amplified enhancers were inserted either directly upstream of the *sna* promoter or downstream of the αtub 3′ UTR. “Enhancer only” and “promoter” only constructs were generated by starting with the pbphi vector and inserting the *sna* shadow enhancer or the *eve* promoter-*PP7-yellow-*αtub 3′ UTR. These constructs were integrated into the fly genome using PhiC31-mediated site specific integration, using the VK00033 and the VK00002 landing sites for the 3rd chromosome and 2nd chromosome insertion, respectively (K. J. T. Venken et al., 2006; Koen J T Venken et al., 2009). Transgenic lines were generated by BestGene Inc.

### Generation of endogenous *sna-MS2* and *sna-PP7* lines

*sna-MS2* and *sna-PP7* fly lines were generated using CRISPR/Cas9-mediated homology-directed repair to insert 24 copies of MS2 and PP7 loops into the 3′UTR of the endogenous *sna* locus. The target site within the 3′UTR was selected using the flyCRISPR Target Finder (Gratz et al., 2014). The following gRNA oligo sequences were used: Sense (5′ - GTC GGG AAT AAT CTT AAC AAC AGT - 3′) and Antisense (5′ - AAA CAC TGT TGT TAA GAT TAT TCC - 3′).

### Generation of homozygous and hemizygous embryos

Males containing the *PP7-yellow* reporter gene were crossed with female *nos* > *MCP-GFP, nos* > *mCherry-PCP, His2Av-eBFP2* flies (Lim, Heist, et al., 2018). The progeny contains one copy of the *PP7-yellow* and one copy of *MCP-GFP, mCherry-PCP, His2Av-eBFP2*, and the females were crossed the *MS2-yellow* containing males. 50% of the resulting embryos carry maternal *PP7-yellow* and paternal *MS2-yellow* reporter genes, which are referred to as “homozygotes.” The remaining 50% carry only the paternal *MS2-yellow* reporter gene, and are referred to as “hemizygotes.” All embryos have one maternal copy of the MCP-GFP, mCherry-PCP, and His2Av-eBFP2, which allows visualization of the *MS2-yellow* and *PP7-yellow* transcriptional activity.

### Live imaging

Embryos were grown and collected at 23°C. Embryos were dechorionated using bleach, washed with water, and mounted on a semipermeable membrane (Sarstedt AG & Co) for imaging. Live images were obtained from a Zeiss confocal microscope, LSM 800, with a Plan-Apochromat 40×1.3 NA oil objective. UV, 488, and 561 lasers were used to visualize His2AV-eBFP2, MCP-GFP, and PCP-mCherry, respectively. The same laser settings were used for all images. The 16-bit images were taken with 1.1x zoom in a 145.20μm × 145.20μm × 9.75μm rectangular prism space where 14 Z-stack was obtained with 0.75μm steps. The temporal resolution was 27 seconds per frame.

### Image Analysis and Statistical Analysis

All the live images were processed and analyzed with Fiji and MATLAB (R2018b, MathWorks) (Schindelin et al., 2012). The raw images were maximum intensity Z projected and concatenated into two-dimensional movies. Each nucleus was segmented by masks which were created in MATLAB and then manually corrected in Fiji. The fluorescenct intensity within each nucleus mask at each time frame was extracted by taking the average of the two highest intensity via MATLAB. The histograms of the snapshots in Figure 1C and Movie 1 were adjusted for visualization purposes only. All analyses were performed on raw images.

The p-values in each plot were calculated with the two-sided Wilcoxon rank sum test, which is equivalent to a Mann-Whitney U-test. The error bars in Figure 3D were calculated from 1000 rounds of bootstrapping with subsample sizes of 50% of the active nuclei of each construct shown.

## Acknowledgments

We thank Mike Levine, Stas Shvartsman, and Lim lab members for helpful discussion. We also thank the FlyBase for providing useful information (REF). This work was supported by the National Science Foundation CAREER MCB 2044613 awarded to B.L.

## Competing Interests

We declare no competing interests

## Supplementary Materials

**Figure S1.**
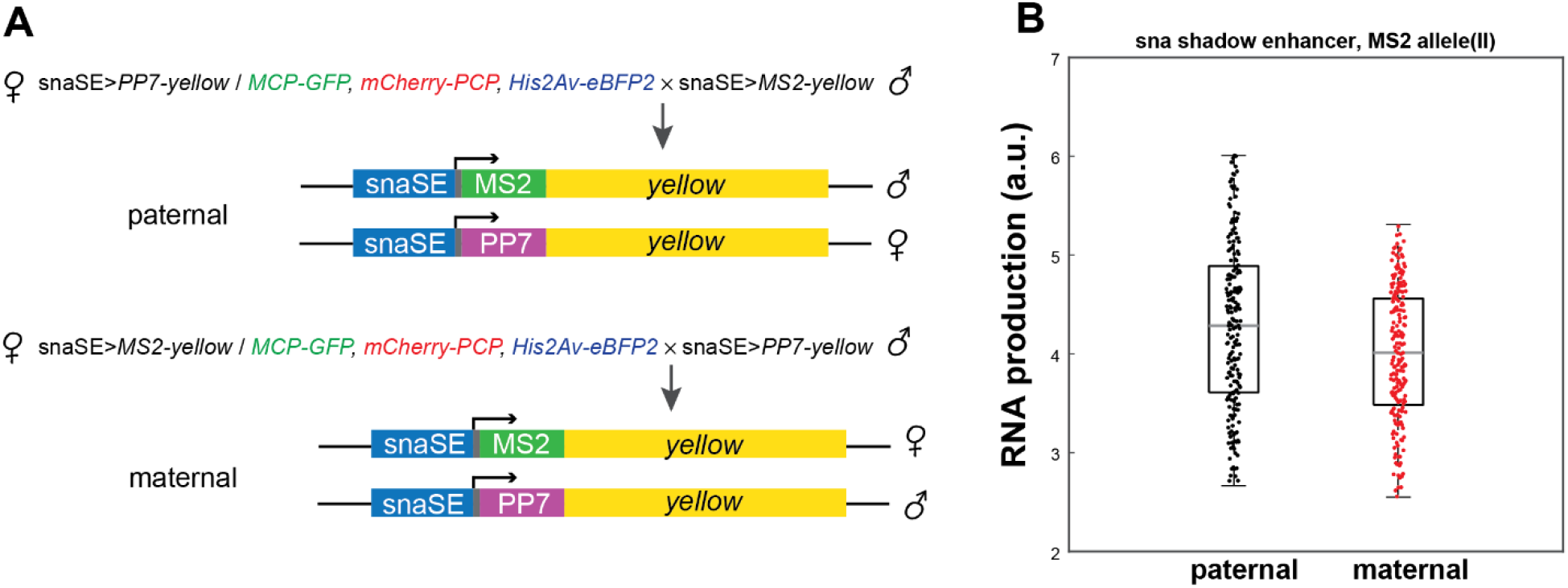
Comparison of the paternal and maternal snaSE>*MS2-yellow* allele. (A) Schematic of genetic crosses to generate embryos with the paternal (up) and the maternal (down) *MS2-yellow* allele. (B) RNA production from paternal and maternal snaSE>*MS2-yellow* allele in snaSE homozygous embryos.

**Figure S2.**
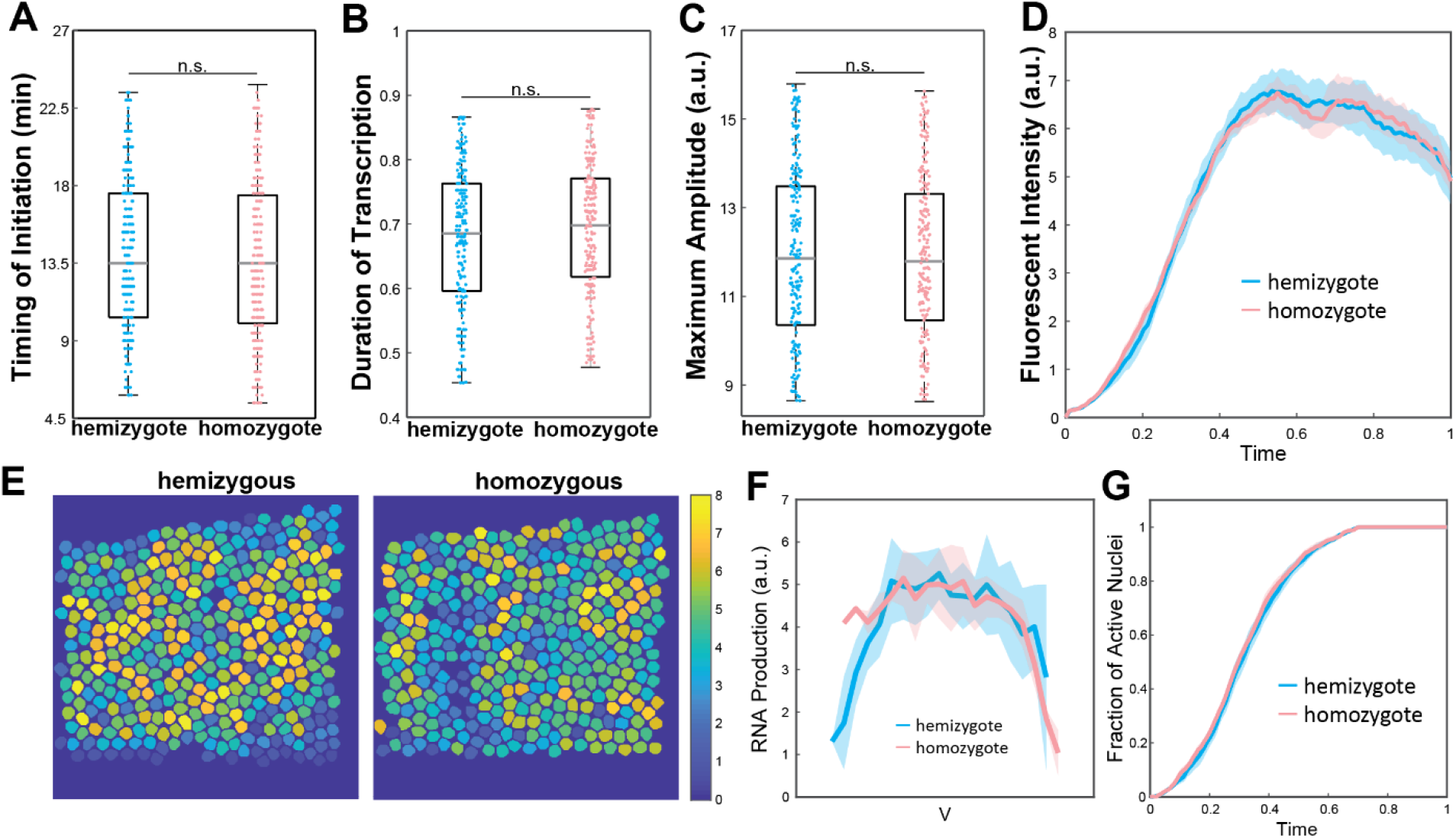
No allelic competition for a weak snaSEmin>*yellow* construct. (A) Boxplot of the timing of transcription initiation for hemizygotes and homozygotes expressing snaSEmin>*yellow*. For both genotypes, transcription was initiated about 13.5 min after the onset of NC14. (B) Boxplot of the duration of active snaSEmin>*MS2-yellow* transcription in NC14. The hemizygous and the homozygous alleles spend comparable time in the active transcription state. (C) Boxplot of the average amplitude of *MS2-yellow* fluorescenct intensity in snaSEmin>*yellow* hemizygotes and homozygotes. The amplitude in homozygous embryos is similar to the one in hemizygous embryos. (D) Average transcriptional trajectories of active nuclei from hemizygous (blue) and homozygous (red) embryos. Two trajectories are mostly overlapping with each other. (E) Heat maps of a representative snaSEmin>*yellow* hemizygous (left) and homozygous (right) embryo showing the accumulated RNA production in NC14 of all nuclei within the *sna* expression domain. The snapshot shows a ventral view of an embryo (F) Average RNA production of hemizygotes (blue) and homozygotes (red) expressing the snaSEmin>*MS2-yellow* reporter gene along the dorsoventral axis of an embryo. The RNA production is comparable throughout the domain and the *sna* expression boundary is not narrowed. (G) Plot of the cumulative fraction of active nuclei over the duration of NC14 in hemizygotes (blue) and homozygotes (red). Both genotypes produce RNAs with similar kinetics of transcriptional activation.

**Movie S1.** Live imaging of hemizygous and homozygous snaSE>*yellow* embryos.

## Notes

### Competing Interest Statement

The authors have declared no competing interest.

